# RRM domain of ALS/FTD-causing FUS interacts with membrane: an anchor of membraneless organelles to membranes?

**DOI:** 10.1101/122671

**Authors:** Yimei Lu, Liangzhong Lim, Jianxing Song

**Affiliations:** Department of Biological Sciences, Faculty of Science, National University of Singapore; 10 Kent Ridge Crescent, Singapore 119260

**Author notes:** The authors have declared that no competing interests exist.

**Keywords:** ALS, FTD, FUS, Low complexity (LC) domain, Membraneless organelle, RNA-recognition motif (RRM), Membrane, NMR spectroscopy

## Abstract

526-residue FUS functions to self-assemble into reversible droplets/hydrogels, which could be further solidified into pathological fibrils. FUS is composed of N-terminal low-sequence complexity (LC); RNA-recognition motif (RRM) and C-terminal LC domains. FUS belongs to an emerging category of proteins which are capable of forming membraneless organelles in cells via phase separation. On the other hand, eukaryotic cells contain a large network of internal membrane systems. Therefore, it is of fundamental importance to address whether membraneless organelles can interact with membranes. Here we attempted to explore this by NMR HSQC titrations of three FUS domains with gradual addition of DMPC/DHPC bicelle, which mimics the bilayer membrane. We found that both N- and C-terminal LC domains showed no significant interaction with bicelle, but its well-folded RRM domain does dynamically interact with bicelle with an interface opposite to that for binding nucleic acids including RNA and ssDNA. If this *in vitro* observation also occurs in cells, to interact with membrane might represent a mechanism for dynamically organizing membraneless organelles to membranes to facilitate their physiological functions.

---

Fused in Sarcoma/Translocated in Sarcoma (FUS) is encoded by a gene which was first identified as a fusion oncogene in human liposarcomas^1,2^. The 526-residue FUS protein belongs to the FET protein family, which also includes Ewing RNA binding protein (EWS), and TATA-binding protein associated factor (encoded by TAF15)^3,4^. Increasing evidence suggests that FUS is involved in various cellular processes, including cell proliferation, DNA repair, transcription regulation, and multiple levels of RNA and microRNA processing^5-7^. On the other hand, FUS is extensively involved in the pathology of neurodegenerative diseases, particularly in amyotrophic lateral sclerosis (ALS), frontotemporal dementia (FTD)^3-15^. These results suggest that FUS might have a general role in neurodegenerative diseases.

FUS is intrinsically prone to aggregation^5-10^, which is composed of an N-terminal low-sequence complexity (LC) domain (1-267); an RNA-recognition motif (RRM: 285-371) capable of binding a large array of RNA and DNA^1,16-19^; and C-terminal LC domain (371-526) (Fig. 1A). RRM is one of the most abundant protein domains in eukaryotes, carrying the conserved RNP1 and RNP2 sequence stretches^18^. Previously the RRM domain of FUS has been determined by NMR spectroscopy to adopt the same overall fold as other RRMs, which consists of a four-stranded β-sheet and two perpendicular α-helices. Very amazingly, although no ALS-causing mutation was identified within RRM, *in vivo* studies revealed that RRM is required for manifesting FUS cytotoxicity ^20^.

**Fig. 1.**
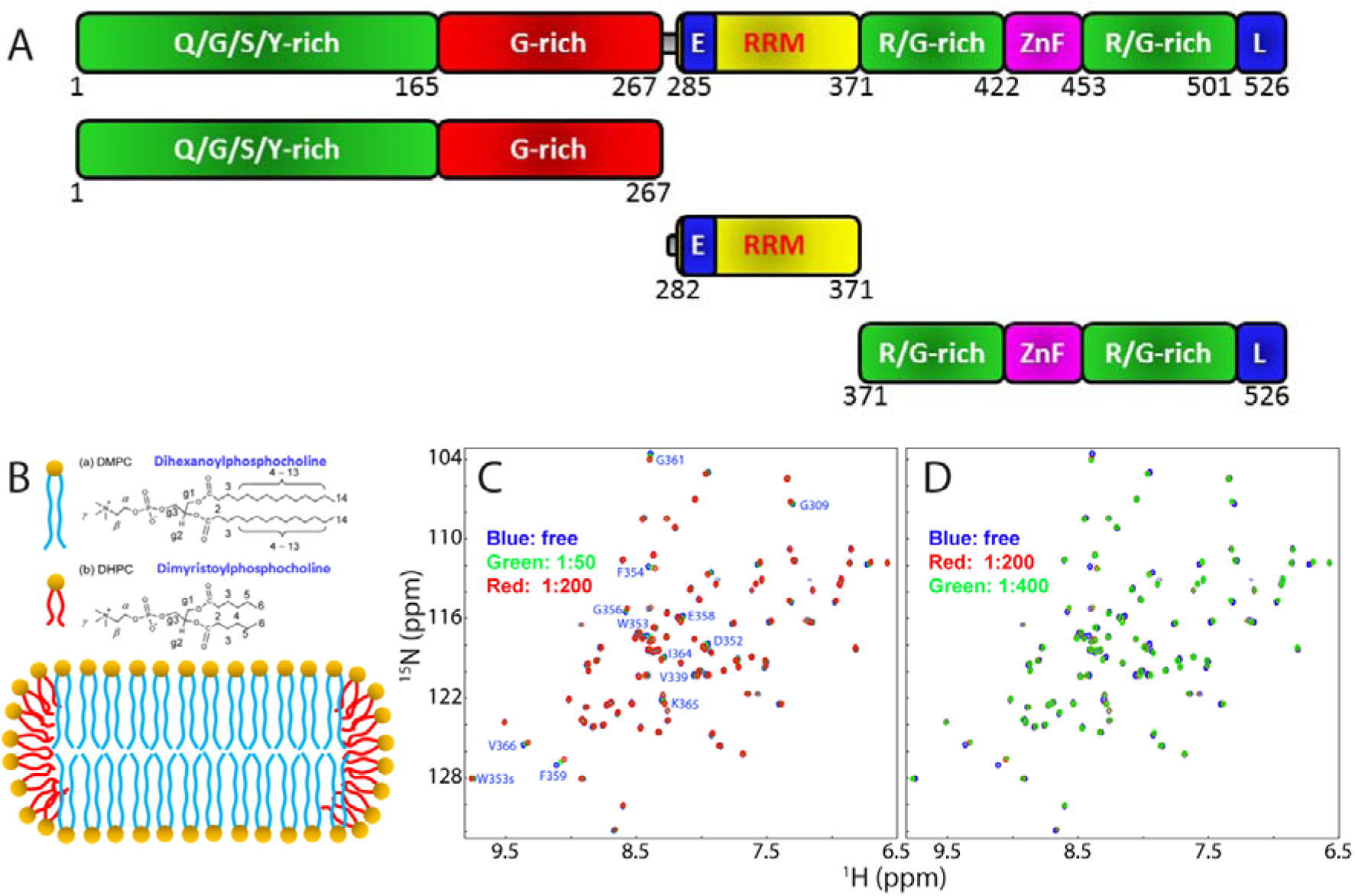
FUS-RRM interacts with membrane. (A) Domain organization of the 526-residue FUS protein; and its three dissected domains. (B)Chemical structures of DMPC and DHPC; as well as bicelle structure formed by DMPC/DHPC. (C)-(D) Superimposition of the two-dimensional NMR ^1^H-^15^N HSQC spectra of the ^15^N-labeled FUS-RRM at a concentration of 100 μM in 10 mM sodium phosphate at pH 6.8 in the absence (blue), and in the presence of bicelle at different ratios.

Previously, as facilitated by our discovery that unlike well-folded proteins following the “Salting-in” rule that protein solubility increases upon adding salts over the range of low salt concentrations (usually < 300-500 mM), “insoluble” proteins could only be solubilized in aqueous solution with minimized salt concentrations^21-23^, we have successfully studied the ALS-causing and aggregation-prone TDP-43 N-terminal and C-terminal prion-like domains^24,25^. The results decoded that there exists a membrane-interacting subdomain flanked by two prion-like sequences within the TDP-43 C-terminal domain (CTD) ^25^. Interestingly, we found that ALS-causing genetic, pathological or environmental factors act to eliminate the native folds of human VAPB-MSP domain and superoxide dismutase 1 (SOD1) and consequently the mutants become predominantly disordered in solution without any stable secondary and tertiary structures, which thus have their intrinsic hydrophobic/amphiphilic regions unavoidably exposed to bulk solvent^26-30^. Therefore, these disordered mutants are only soluble in unsalted water but got aggregated in high salt solutions such as in cells with ~150 mM salts. Most unexpectedly, they acquired novel capacity in interacting with membranes energetically driven by forming helices in membrane environments^21-23,26-30^.

Here, we attempted to explore whether FUS domains are able to interact with membranes by monitoring the peak shift of their HSQC peaks upon gradual addition of DMPC/DHPC bicelle, which mimics the bilayer membrane (Fig. 1B). Interestingly, we found that FUS LC domains showed no significant interaction with DMPC/DHPC bicelle, but its well-folded RRM does dynamically interact with bicelle with an interface opposite to that used for binding nucleic acids including RNA and ssDNA as previously mapped out^17^.

## Results

We first cloned and expressed three FUS domains: namely the N-terminal LC (1-267), RRM (282-371) and C-terminal LC (371-526) domains. While the two LC domains were all found in inclusion body and thus purified under denaturing conditions; the FUS-RRM domain is highly soluble in supernatant and purified under native conditions. Subsequently, we acquired series of HSQC spectra of the ^15^N-labeled FUS domains in the presence of DMPC/DHPC bicelle (Fig. 1B) at different ratios. The N- and C-terminal LC domains of FUS showed no significant shift of their HSQC peaks even in the presence of bicelle at a ratio up to 1:800 (FUS domain:bicelle), implying that they have no significant ability to interact with membranes.

Unexpectedly, however, upon stepwise addition of bicelle, a subset of HSQC peaks of the RRM domain underwent gradual shifts (Fig. 1C). Furthermore, the shifts were mostly saturated at ratio of 1:200 (Fig. 1D). In order to identify the FUS residues with shifted HSQC peaks, we achieved the sequential assignment of our RRM (282-371) by analyzing a pair of triple resonance spectra HN(CO)CACB and CBCA(CO)NH collected on a ^15^N-/^13^C-double labeled sample. Fig. 2A presents the (∆Cα−∆Cβ) chemical shifts, which represent a sensitive indicator of the secondary structures of proteins^31^. In addition to the N-terminal residue Ser282, as well as Pro320, Pro344, Pro345 and Pro363, only Lys315, Glu336, Asp343 and Ser360 were not assigned due to overlap or undetectable resonance signals. Most (∆Cα−∆Cβ) chemical shifts of our FUS RRM construct are almost the same as those associated with the NMR structure of FUS-RRM (BMRB ID of 17635) over the identical region^17^.

**Fig. 2.**
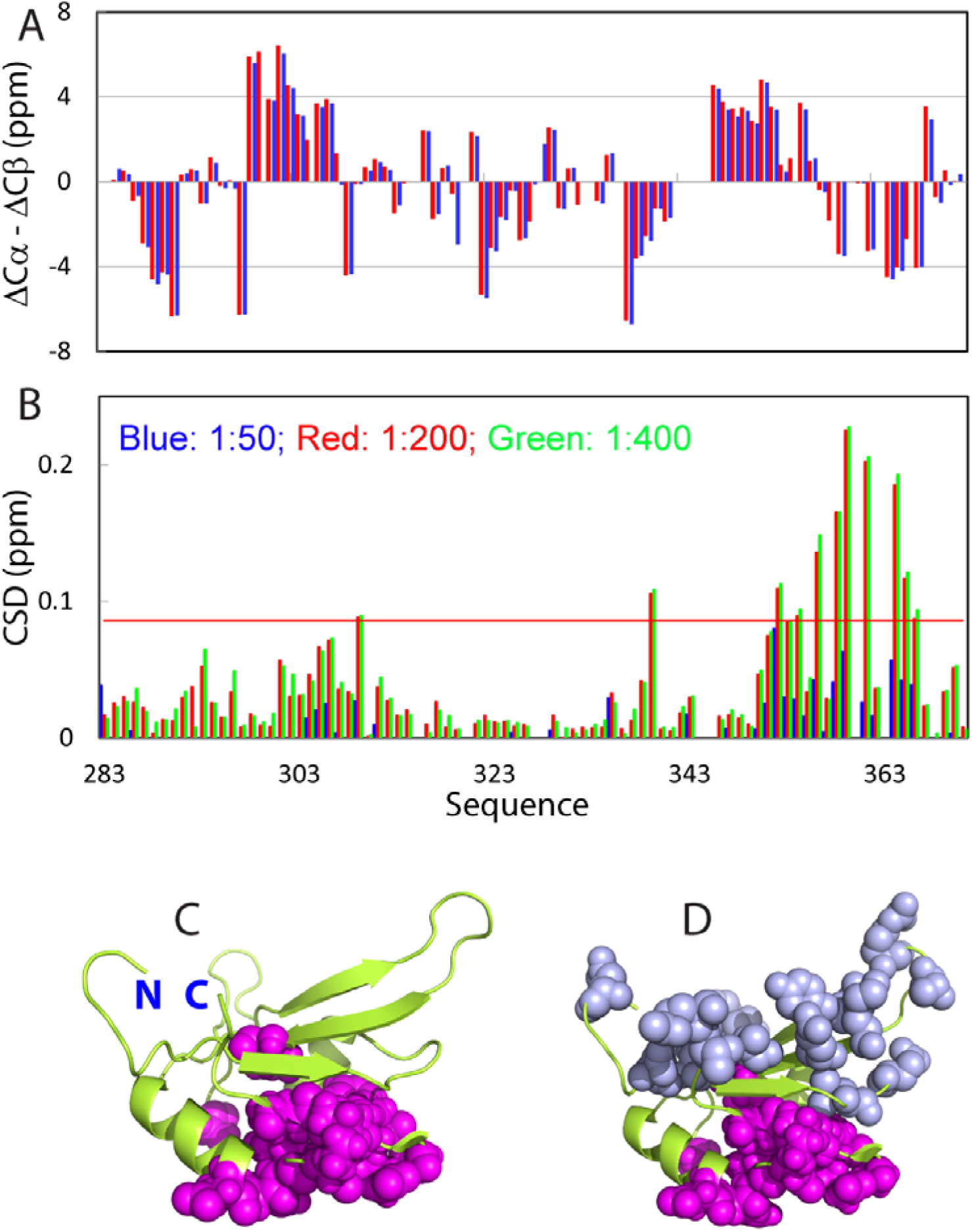
Interface of FUS-RRM for interacting with membrane. (A) Residue specific (∆Cα−∆Cβ) chemical shifts of the FUS RRM domain collected at 25 °C in 10 mM phosphate buffer at pH 6.8 (blue) and those previously deposited in BMRB (ID of 17635) (red). (B) Chemical shift difference (CSD) of FUS-RRM in the presence of bicelle at different ratios (RRM:bicelle): 1:50 (blue); 1:200 (red) and 1:400 (green). (C) NMR structure of FUS-RRM (PDB ID of 2LCW) with the residues having significant shifts of HSQC peaks (> average + STDEV [0.086 ppm]) displayed in purple spheres. (D) NMR structure of FUSRRM (PDB ID of 2LCW) further with residues involved in binding ssDNA previously reported^17^ displayed in light blue spheres.

Fig. 2B shows the residue-specific chemical shift difference (CSD) of FUS-RRM in the presence of bicelle at different ratios. Interestingly, upon mapping back the residues with significant HSQC peak shifts to the NMR structure, an interesting picture emerges (Fig. 2C): these residues are clustered together to form an interface located on one side of the structure.

We also titrated our FUS-RRM domain with a single-strand DNA (ssDNA) previously used; and found that the residues with significant shifts upon adding this ssDNA are identical to those previously mapped out^17^. Strikingly, the set of residues involved in binding ssDNA have no overlap with the residues whose HSQC peaks significantly shifted upon adding bicelle. The residues critical for binding nucleic acids^17^ are located on the opposite side of the FUS-RRM structure (Fig. 2D).

## Discussion

Recently, it has been deciphered that proteins containing the LC domains such as prion-like sequences are significantly over-represented by RNA-/DNA binding proteins as exemplified by FUS and TDP-43^32-44^. Strikingly, these proteins have been increasingly characterized to be involved in various diseases, particularly neurodegenerative diseases. Most remarkably, their prion-like domains have unique capacity in self-assembling to achieve phase separation which can further transform into fibrils with cross-β structures. While it remains highly controversial what are the roles of the diverse structures formed during self-assembly in physiology and pathology, it becomes clear that the phase separation represents a key mechanism to form membraneless compartments or organelles in cells^32-51^.

On the other hand, eukaryotic cells contain a large network of internal membrane systems. This inspires us to hypothesize that the cellular functions of some membraneless organelles formed by phase separation might be facilitated if they can have interactions with membranes to different degrees. For example, the localization to membrane might increase the local concentration of these proteins to enhance self-assembly. Indeed, previously, we found the presence of a hydrophobic subdomain within the TDP-43 CTD, which can transform from partially folded state into a well-folded Ω-loop-helix motif upon interacting with membranes^25^. Most importantly, we showed that the self-assembly was significantly enhanced if the TDP-43 CTD was anchored into bicelle^25^.

Here we found the well-folded FUS-RRM domain also interacts with bicelle by using a small set of residues. Previously, we have investigated but found that the wild-type VAPB-MSP domain and the matured SOD1 showed no detectable interaction with bicelle and even DPC-micelle^26-28^. Moreover, the interfaces of FUS-RRM for interacting with nucleic acids and membrane are located on opposite sides of the structures. This strongly implies that our current *in vitro* observation that FUS-RRM is capable of interacting with membrane might indeed occur in cells. Nevertheless, this observation certainly needs *in vivo* confirmation although it might be extremely challenging due to its dynamic nature.

## Methods

### Preparation of recombinant proteins

The DNA fragments encoding three dissected domains of FUS (Fig. 1A) were amplified by PCR reactions from FUS cDNA and subsequently cloned into a modified vector pET28a as we previously used for the TDP-43 prion-like domain^25^. The expression vectors were subsequently transformed into and overexpressed in *Escherichia coli* BL21 (DE3) cells (Novagen). The recombinant proteins of FUS (1-267) and FUS (371-526) were found only in inclusion body while RRM was highly soluble in supernatant. As a result, for FUS (1-267) and FUS (371-526), the pellets were first dissolved in a phosphate buffer (pH 8.5) containing 8 M urea and subsequently purified by a Ni^2+^-affinity column (Novagen) under denaturing conditions in the presence of 8 M urea. The fractions containing the recombinant proteins were acidified by adding 10% acetic acid and subsequently purified by reverse-phase (RP) HPLC on a C4 column eluted by water-acetonitrile solvent system. The HPLC elution containing pure recombinant proteins were lyophilized. For RRM, the recombinant proteins were purified by a Ni^2+^-affinity column (Novagen) under native condition, followed by a further purification by FPLC on a gel-filtration column.

The generation of the isotope-labelled proteins for NMR studies followed a similar procedure except that the bacteria were grown in M9 medium with the addition of (^15^NH_4_)_2_SO_4_ for ^15^N labeling and (^15^NH_4_)_2_SO_4_/[^13^C]-glucose for double labelling^24,25^. The purity of the recombinant proteins was checked by SDS–PAGE gels and their molecular weights were verified by a Voyager STR matrix-assisted laser desorption ionization time-of-flight-mass spectrometer (Applied Biosystems). The concentration of protein samples was determined by the UV spectroscopic method in the presence of 8 M urea. Briefly, under the denaturing condition, the extinct coefficient at 280 nm of a protein can be calculated by adding up the contribution of Trp, Tyr and Cys residues^56^.

### NMR experiments

All NMR experiments were acquired on an 800 MHz Bruker Avance spectrometer equipped with pulse field gradient units as described previously^24,25^. For characterizing the solution structure of the FUS RRM domain, a pair of triple-resonance experiments HNCACB, CBCA(CO)NH were collected for the sequential assignment on a ^15^N-/^13^C-double labelled sample of 400 μM.

Here a membrane-mimetic system DMPC/DHPC bicelle was used to assess the membrane-interaction of FUS-RRM. The large bicelle to better mimic bilayer membrane was prepared by mixing up DMPC and DHPC at a q value of 4 as previously described^25,53^. At this ratio, the disk-shaped bicelle with a diameter of ~460 Å is formed in which DMPC constitutes a bilayered section surrounded by a rim of DHPC (Fig. 1B). The FUS-RRM sample for titrating with bicelle was prepared at 100 μM in 10 mM sodium phosphate buffer at pH 6.8.

## Acknowledgement

This study is supported by Ministry of Education of Singapore (MOE) Tier 2 MOE2015-T2-1-111 to Jianxing Song. The funders had no role in study design, data collection and analysis, decision to publish, or preparation of the manuscript.

## Author Contributions

Conceived and designed the experiments: JXS; Performed the experiments: YML LZL. Analyzed the data: YML LZL JXS. Prepared figures and wrote the paper: JXS.

